# ExoJ: an ImageJ2/Fiji plugin for automated spatiotemporal detection and analysis of exocytosis

**DOI:** 10.1101/2022.09.05.506585

**Authors:** Junjun Liu, Frederik Johannes Verweij, Guillaume Van Niel, Thierry Galli, Lydia Danglot, Philippe Bun

**Author notes:** Co-senior authors (Lydia Danglot, Philippe Bun).

## Abstract

Exocytosis is a dynamic physiological process that enables the release of biomolecules to the surrounding environment via the fusion of membrane compartments to the plasma membrane. Understanding its mechanisms is crucial, as defects can compromise essential biological functions. The development of pH-sensitive optical reporters alongside fluorescence microscopy enables the assessment of individual vesicle exocytosis events at the cellular level. Manual annotation represents, however, a time-consuming task, prone to selection biases and human operational errors. Here, we introduce ExoJ, an automated plugin based on ImageJ2/Fiji. ExoJ identifies user-defined genuine populations of exocytic events, recording quantitative features including intensity, apparent size and duration. We designed ExoJ to be fully user-configurable, making it suitable to study distinct forms of vesicle exocytosis regardless of the imaging quality. Our plugin demonstrates its capabilities by showcasing distinct exocytic dynamics among tetraspanins and vesicular SNAREs protein reporters. Assessment of performance on synthetic data demonstrated ExoJ is a robust tool, capable to correctly identify exocytosis events independently of signal-to-noise ratio conditions. We propose ExoJ as a standard solution for future comparative and quantitative studies of exocytosis.

## Introduction

Exocytosis is a fundamental biological process to convey sets of chemical information to the extracellular environment, and deliver membrane proteins and lipids to the plasma membrane (PM) (Jahn and Südhof, 1999). Briefly, the process consists in the transport, docking, priming and fusion of intracellular compartments to the PM (Verhage and Sørensen, 2008). This eventually leads to the deposition of proteins and lipids into the PM, and the release of their luminal content to sustain important physiological functions and to respond to external stimuli. An orchestrated spatiotemporal control of exocytosis is thus critical in the regulation of physiological and pathological processes such as those sustained by the release of exosomes (Alberts et al., 2006; Bebelman et al., 2018; Chen and Scheller, 2001; Guček et al., 2019; Gundelfinger et al., 2003; Lamichhane et al., 2015; Martinez-Arca et al., 2001a; Meldolesi, 2018; Urbina et al., 2018; van Niel et al., 2018; Wang et al., 2018). This also underlies the need to characterize the structural dynamics of the fusion machinery, the biochemical profile of the biological contents and the spatiotemporal dynamics of their release. In addition to a collection of biochemical approaches, methods using live-cell imaging of fluorescently-tagged intracellular vesicles allow for spatiotemporal monitoring and quantitative analysis of content release (Ge et al., 2010). In particular, total internal reflection fluorescence microscopy (TIRFM) has become the imaging modality of choice due to its inherent high signal-to-noise ratio with a reduced phototoxicity while increasing temporal resolution (Axelrod and Omann, 2006; Bebelman et al., 2020; Burchfield et al., 2010; Mattheyses et al., 2010; Miesenböck et al., 1998; Sankaranarayanan et al., 2000a; Schmoranzer et al., 2000). The TIRFM evanescent field of illumination enables the recording of events close to the PM, minimizing the disturbance from fluorescently-labeled vesicles in the cytoplasm. Specific labeling of vesicle exocytosis is achieved by tagging the content and/or the vesicle membrane with a pH-sensitive fluorescent protein (FP) variant of GFP, known as pHluorin (Miesenböck et al., 1998). The FP pHluorin is quenched in the acidic vesicular lumen, and brightens in the neutral extracellular environment upon vesicle fusion to the PM (hereafter termed fusion event). Using TIRFM, the fusion event appears as an abrupt brightening followed by spreading of the fluorescence signal (Miesenböck et al., 1998). Besides the vesicle-specific protein marker, the choice of the FP greatly influences the monitoring of the dynamics of exocytosis steps as evidenced by others (Liu et al., 2021a; Martineau et al., 2017; Sankaranarayanan et al., 2000b; Shen et al., 2014). Taken together, the subsequent amount of data generated by fluorescent time series poses the need of a robust analysis pipeline to identify exocytic events in an unbiased manner. Manual annotation of each candidate event represents a time-consuming task, and is prone to selection biases. In particular, recordings of fusion event XY location to report potential intracellular hotspots, the intensity which estimates the relative amount of proteins, the apparent size and the duration are hardly reproducible from one cell to the other. Numerous algorithms have been developed to address these challenges in a semi-(Bebelman et al., 2020; Huang et al., 2007; Jullie et al., 2014; Wang et al., 2018) or fully automated manner (Diaz et al., 2010; Mahmood et al., 2023; Moro et al., 2021a; Persoon et al., 2019; Sebastian et al., 2006; Urbina and Gupton, 2021; Yuan et al., 2015). These solutions are, however, optimized for a particular population of vesicular cargoes and hence specific applications (Bebelman et al., 2020; Burchfield et al., 2010; Diaz et al., 2010; Huang et al., 2007; Jullie et al., 2014; Moro et al., 2021b; Persoon et al., 2019; Sebastian et al., 2006; Urbina et al., 2018; Wang et al., 2018; Yuan et al., 2015). Here we present Project-ExoJ (hereafter named ExoJ), a solution developed as an ImageJ2/Fiji (Schindelin et al., 2012) plugin that automates the identification of fluorescently-reported fusion event. We designed ExoJ to be fully user-configurable, with a graphical user interface to set up a series of parameters that define a genuine population of exocytic events according to the experimental conditions. To improve user experience, we provided tools for visualizing and further reporting features such as spatial location, intensity over time, apparent size and duration. To illustrate ExoJ capability, we focused on fusions events reported by tetraspanin (TSPAN) and vesicular soluble N-ethylmaleimide-sensitive fusion protein attachment protein receptor (v-SNARE) proteins coupled to pHluorin. Recordings of quantitative features revealed significant differences between and among labeled vesicle populations. Assessment of ExoJ performance using synthetic data revealed a highly robust and reliable identification tool insensitive to noise encountered in experimental settings.

## Results

### ExoJ workflow for automated identification of exocytic events

At least three modes of exocytosis have been identified (full-collapse, kiss-and-run and compound exocytosis) and characterized according to their fusion dynamic patterns (Wu et al., 2014). Although each mode has its own fluorescence fluctuation pattern, the intensity decays to a certain extent right after vesicle fusion to the PM. This makes automatic recognition possible. Hence, we describe a fusion event as a transient diffraction-limited or large object that displays a sudden increase followed by an exponentially decreasing fluorescence intensity into the background. This fluorescence decrease is characterized by the mean lifetime decay τ (Fig. 1A). With this definition, we designed a processing pipeline broken into three main steps. Each step is managed in its own graphical user interface (GUI) dialog in which every parameter can be manually adjusted (Fig. 1). These parameters can be saved and called in the pipeline to improve reproducibility (Fig. 1B,C,D and Table 1).

**Figure 1.**
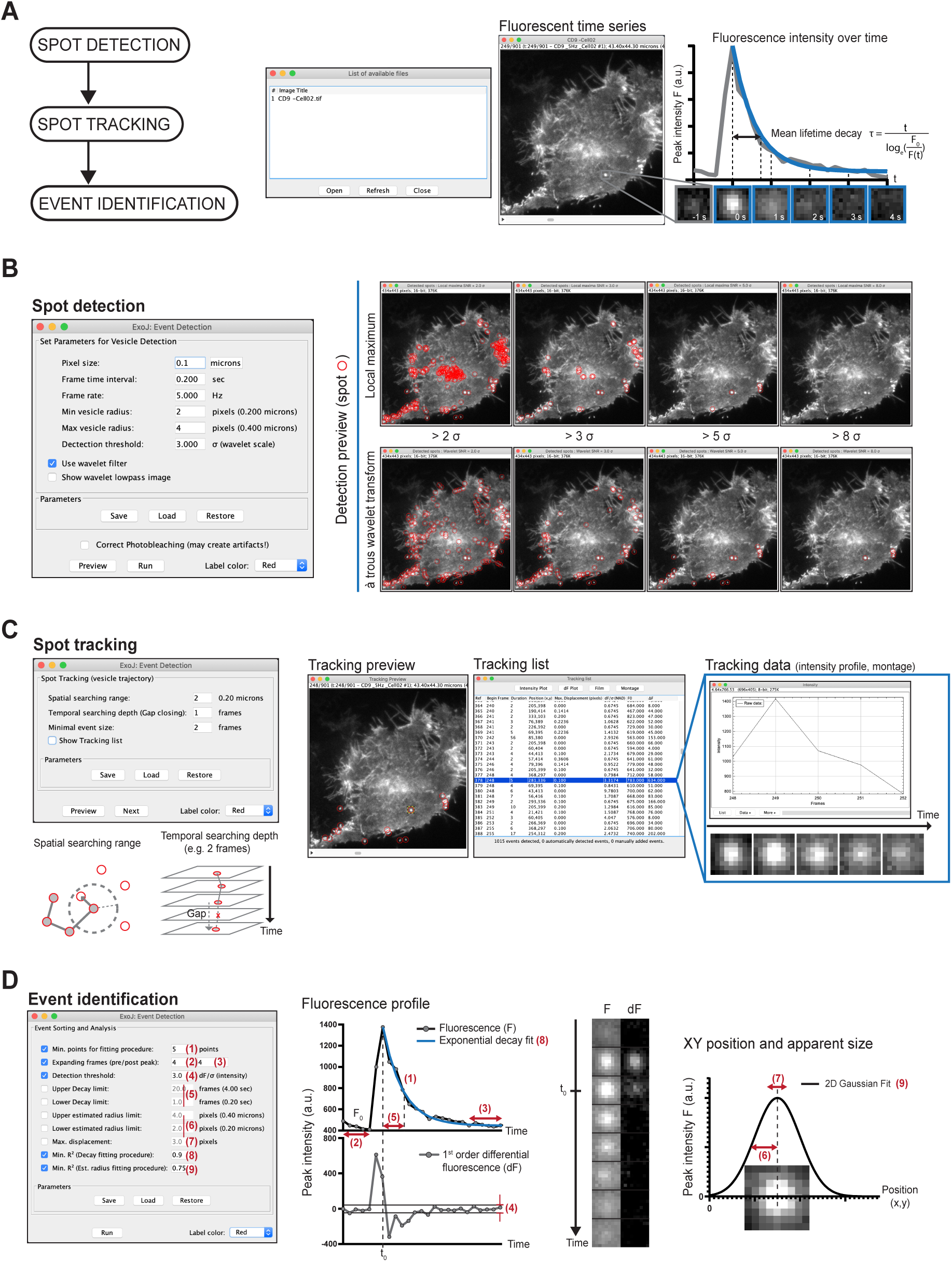
ExoJ facilitates the automated identification of pHluorin-reported exocytosis. **(A)** A prompt of available files appears at the start of ExoJ. Users can load, refresh and select image series of interest. The framework of the algorithm comprises three main steps to identify and analyze exocytic events from fluorescent time series. A typical fusion event labeled by CD9-pHluorin in Hela cell is shown as a time-lapse montage. Before fusion (gray), the fluorescence is quenched in acidic environment. Upon fusion, the pHluorin is exposed to neutral pH. A bright diffraction-limited spot appears and diffuses over time (blue-colored inset frames). Signal intensity is fitted using a mono-exponential decay function to extract the mean lifetime, denoted as τ. **(B)** Detection of vesicles (seens as spots) are first defined by setting the minimal and maximal vesicle radius. Vesicles are detected using either a local maximum (unchecked box) or à trous wavelet transform algorithm (default option, checked box). The detection threshold value is manually set by users to permit the algorithm to detect spots on single images (here in red). The threshold value is a multiple of σ (MAD) of either wavelet coefficients C_wavelet_ (à trous wavelet transform) or pixel intensity calculated for each fluorescent image (local maximum). Increasing the detection threshold ultimately leads to a decreased number of detected vesicles as illustrated for both algorithms here. **(C)** Previously detected spots are connected to reconstruct time-lapse trajectories. The parameters considered for reconstruction are illustrated, and include the maximal distance (spatial searching range) and the time gap (temporal searching depth) between two successive spots as well as the minimal number of connected spots (event duration). Tracking results including intensity profile and fluorescent montage for single candidate events can be reviewed upon selection (Show Tracking list button). **(D)** For each candidate fusion event, a series of user-defined parameters are applied to identify fusion events. (1) The number of data points used for fitting a negative mono-exponential function to derive the mean lifetime τ can be adjusted. (2-3) The number of additional frames around the detected event (intensity peak) is set by users to refine the evaluation of local background. (4) The detection threshold σ_dF_ is the MAD of the 1^st^ order differential fluorescence intensity profile set to sort exocytic events. (5) Upper and lower decay limit entries enable users to set the range of mean lifetime τ of fusion events. The fluorescence peak intensity profiles at different timepoints were fitted with a two-dimensional Gaussian fit to estimate the apparent radius of fusion event and track the XY position. Fusion events can be identified by accordingly setting the range of apparent radius (6) and the position XY over time (7). Evaluation of both mean lifetime τ and apparent radius is associated with the goodness-of-fit R^2^ (8-9).

**Table 1.** General commands throughout the identification and analysis process.

|  |  |
| --- | --- |
| **(Pop-up window)**<br>**List of available files** | Upon starting the plugin, a pop-up window lists opened image-type files. To start (resp. stop) the analysis, press **Open** (resp. **Close**). If an image series does not appear, **Refresh** the pop-up window.<br>After pressing **Open**, the selected image series will be duplicated for further processing and analysis. Note that the plugin can run in parallel multiple instances. |
| **Save, Load, Restore** | Parameters are saved (in .dat format), loaded (.dat file) or restored to the metadata of the image series. |
| **Label color** | Detected vesicles (seen as spots) are marked by circles of selected color. |
| **Preview, Run** | At each step, processing results can be reviewed for different set of parameters (**Preview**).<br>During the spot detection step, single fluorescent images are displayed. During the spot tracking step, the whole sequence is shown. During the event identification step, the whole sequence and the table result are displayed.<br>To proceed to the next step, press **Run**. |

#### Vesicle detection

The first step consists in detecting vesicles seen as bright spots from image series (Fig. 1B and Table 2). Before performing spot detection, we apply a custom photobleaching correction algorithm to compensate for the variations in image intensity within fluorescent time sequences, as described in the Materials and Methods section. Users can rely on either a wavelet-based (default option) or local maximum algorithm detection (by unchecking the wavelet filter box; Fig. 1B). For the wavelet-based option, we employ the multiscale à trous wavelet transform algorithm (Olivo-Marin, 2002).

**Table 2.** Parameters and options for vesicle (seen as spot) detection.

|  |  |  |
| --- | --- | --- |
| **Pixel size, Frame rate, Frame time interval** |  | Pixel size, frame rate (and time interval) are automatically extracted from the metadata of selected image series. Correct if necessary. |
| **Min. and Max. vesicle radius** |  | Initial user-entries for the estimated radius (in pixel) of the vesicular population of interest. |
| **(Checked)**<br>**Use wavelet filter** | | Apply à trous wavelet transform algorithm on fluorescent image series where pixel values now correspond to wavelet coefficient ($C_{\text{wavelet}}$). Result on single images can be reviewed (Preview button). |
| | **Detection threshold** | Critical integer value that corresponds to a multiple of the MAD of $C_{\text{wavelet}}$ (noted $\sigma_{\text{wavelet}}$). For a high (resp. low) value, number of detected spots is decreasing (resp. increasing) along with the calculation time. |
| | **Show lowpass image** | When activated, this option displays the result of the à trous wavelet transform algorithm on single frames including fluorescent spots (colored circles) with $C_{\text{wavelet}}$ above the **detection threshold**. |
| **(Unchecked)**<br>**Use wavelet filter** |  | The spot detection relies on the local maximum algorithm with **Min.** and **Max. vesicle radius** as initial entries. |
| | **Detection threshold** | Critical integer value that corresponds to a multiple of the MAD of all pixel intensity (noted $\sigma_{\text{intensity}}$) at each image. |
| **Correct Photobleaching** |  | A pixel-by-pixel detrending method is immediately applied to the image series. Strongly recommend to correct photobleaching when applicable to improve wavelet-based spot detection. Note that the correction can introduce bright/saturated pixels. The process can be reversed by deactivating the option. |

To selectively keep user-specified sized spots from image series, images are convolved with a set of wavelet functions that are scaled according to the minimum and maximum vesicle radius (Min. and Max. vesicle radius; Fig. 1B). The resulting images are then decomposed into high and low-frequency components. On the low frequency component images, pixel values are now equal to wavelet coefficients C_wavelet_ as a result of wavelet transformation (Show wavelet lowpass image; Fig. 1B). The coefficient C_wavelet_ coarsely translates the similarity between sets of pixels and the user-defined wavelet functions. C_wavelet_ value increases when there is a close resemblance between the intensity signal and the user-defined wavelet functions.

The median absolute deviation (MAD) is a measure of variability similar to standard deviation but less sensitive to outliers (e.g. non-specific cell compartments, noise saturated pixels, …). We next define a single hard threshold parameter kσ_wavelet_ with σ_wavelet_ calculated from the MAD of C_wavelet_ as follows:

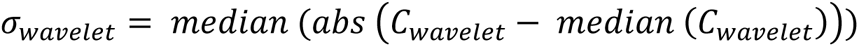

Users set the tolerance of the spot detection algorithm when they set the value of k. Thus, the algorithm sets to zero the pixel coefficients C_wavelet_ whose absolute values are lower than kσ_wavelet_ (Fig. 1B). Hence, setting a high k value would decrease the number of detected objects. An inverse transform is finally applied to reconstruct single fluorescence intensity images (Fig. 1B and Table 2). Alternatively, users can rely on a custom-written local maximum algorithm on single images (unchecked Wavelet Filter box; Fig. 1B). In this option, each pixel is replaced with its corresponding neighborhood maximum intensity value. The radius of the neighborhood is set to twice the user-defined minimal vesicle size. With this approach we also defined kσ_F_ where σ_F_ is now obtained from the MAD of pixel intensity at each image.

The detection result on single images can be previewed to ensure that proper fluorescence spots were accurately detected (Fig. 1B and Table 2). Once determined, the parameter is applied for the whole image series to extract the XY location of all fluorescent spots with a local maxima method within an adaptive window sized to 2 ∗ *Min*. *vesicle radius* + 1.

#### Vesicle tracking

The second step consists in building individual time-lapse trajectories of previously localized fluorescent spots (Fig. 1C). While the starts of single trajectories are due to the appearance of a bright spot, the ends are not solely due to the fusion to the PM but could result from limitations in imaging conditions (e.g. low signal-to-noise ratio, loss of focus, …). To account for this caveat, we combine a simple but yet sufficient multi-frame nearest-neighbor approach with a gap-closing algorithm (Chenouard et al., 2014; Crocker and Grier, 1996). Here, our plugin introduces three cut-off parameters that need to be tailored according to vesicle behavior, consisting of a spatial searching range, a temporal searching window and a minimal event size (Fig. 1C and Table 3). Spot size, direction and intensity are not considered during the frame-to-frame tracking process. Spots within the user-defined spatial and temporal searching range are assigned to the same trajectory, minimizing their global lateral displacement.

**Table 3.**
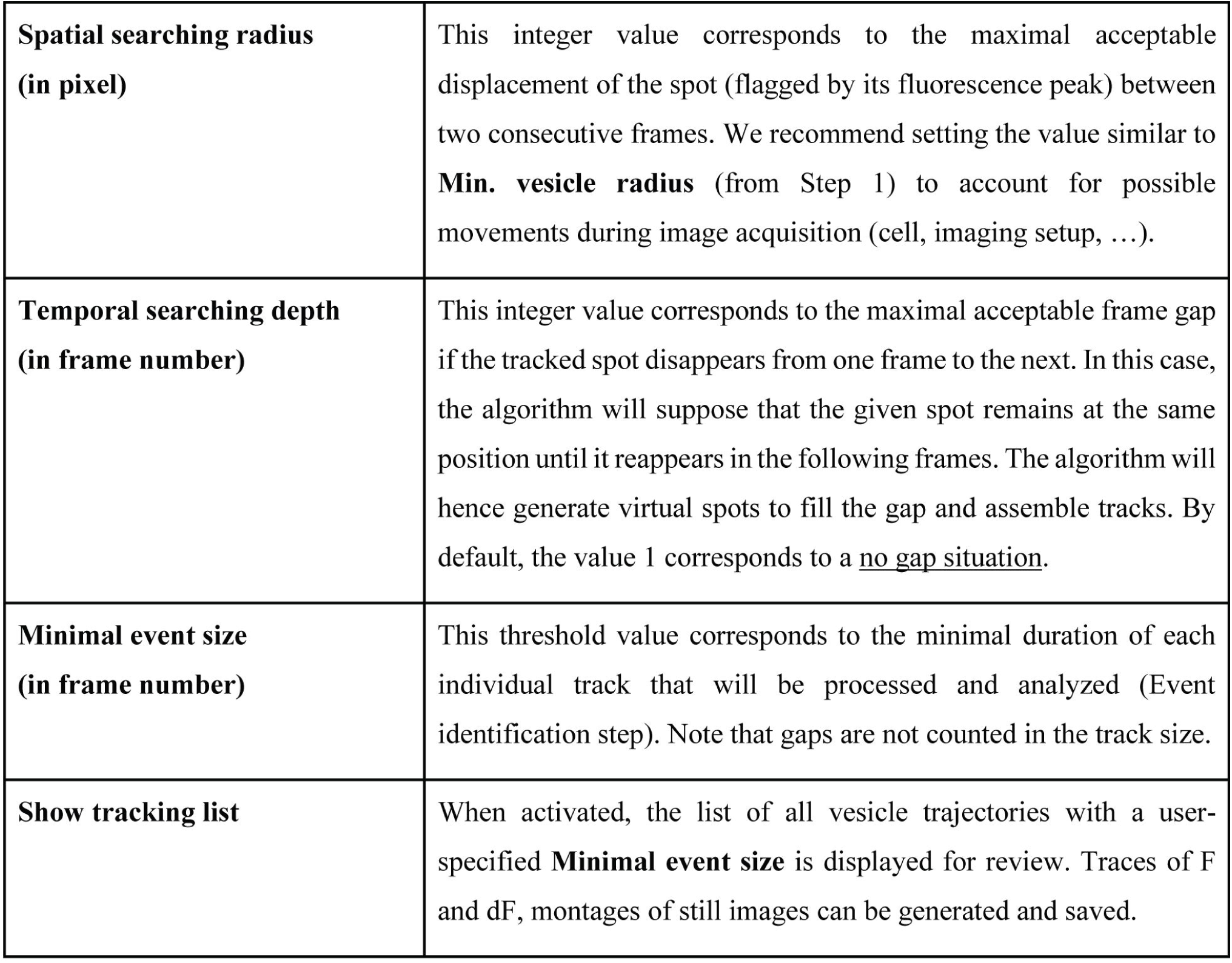
Parameters and options for vesicle (seen as spot) tracking.

#### Fusion event identification

Various features have been considered to streamline the accurate identification of different types of fusion events in different cellular contexts (Bebelman et al., 2020; Burchfield et al., 2010; Diaz et al., 2010; Moro et al., 2021a; Sebastian et al., 2006; Urbina et al., 2018; Wang et al., 2018; Yuan et al., 2015). Here, we combine advantages of previous methods to define a versatile processing protocol. We reason that all fusion events display a statistically significant and transient fluorescence peak fluctuation F above the local background F_0_, followed by an exponential fluorescence decay (Fig. 1D). Considering single vesicle trajectories, we first perform consecutive adjacent image subtraction to normalize the background fluorescence intensity (Jullie et al., 2014; Sebastian et al., 2006). This step results in a high signal-to-noise ratio 1^st^-order differential fluorescent image series dF (Fig. 1D). For each candidate fusion event, we calculate σ_dF_ as the MAD of dF instead of solely considering the normalized peak change in fluorescence intensity ΔF/F_0_ (Bebelman et al., 2020; Jullie et al., 2014; Persoon et al., 2019; Urbina et al., 2018). To refine the estimation of σ_dF_, the trajectories of single candidate events are extended before (resp. after) the appearance (resp. disappearance) of the vesicle according to user entries (Expanding Frames; Fig. 1D). We eventually proceed with a moving linear regression on the fluorescence peak intensity profile to refine the onset time t_0_ of candidate fusion events. This step helps considering fluorescence saturation and successive events at the same XY location. The algorithm also evaluates the maximal displacement of candidate events relative to their initial XY location at t_0_.

Additional measurements are made by our plugin to describe the population of candidate fusing vesicles, including the mean lifetime τ which relates to the fusion dynamics and serves as a proxy for the fusion duration; the apparent size of the fusion event and the normalized peak change in fluorescence intensity ΔF/F_0_ which estimates the relative amount of fluorescently-labeled proteins.

To account for various types of vesicle fusion and/or image series acquired under different experimental conditions, we integrate user-defined entries to modulate the definition of a genuine fusion event and hence the identification requirements of the algorithm (Table 4). In particular, candidate events with dF higher than kσ_dF_ (Detection threshold; Fig. 1D), limited displacements (Max. displacement), duration (Upper/Lower decay limit) and estimated size (Upper/Lower est. radius limit) at t_0_ that comply with user inputs are seen as genuine fusion events (Fig. 1D). For the last two parameters, the goodness-of-fit, reported by the coefficient of determination R^2^, is set as a threshold value (Min. R^2^) above which events are selectively kept (Fig. 1D, see parameters 8 and 9). Once the identification parameters are set, the plugin summarizes features of user-defined genuine events for potential manual curation, visualization and export of detection features (Fig. S1).

**Table 4.**
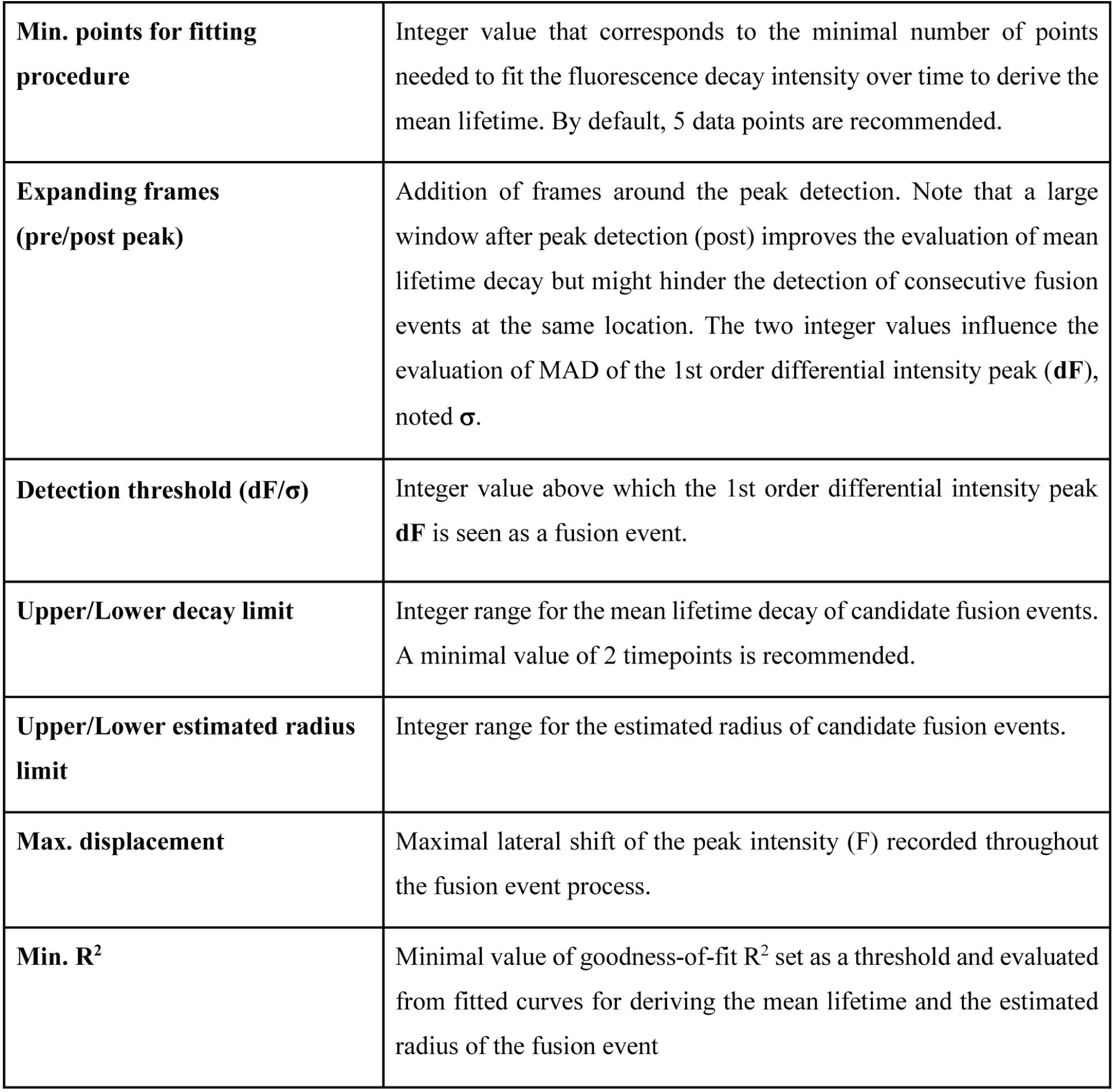
Parameters and options for the identification of vesicle fusion events.

### ExoJ is an adaptative tool to detect fusion events with high accuracy

To evaluate whether our tool can accurately detect fusion events, we used Hela cells expressing pHluorin-tagged CD9, CD63 and CD81 TSPANs commonly used to track extracellular vesicle (EV) exocytosis (Crescitelli et al., 2013; Kowal et al., 2016; Mathieu et al., 2021; Théry et al., 2018; Witwer and Théry, 2019), and v-SNARE VAMP2 and VAMP7 proteins which are components in the fusion machineries in many cellular contexts (Burgo et al., 2012; Chaineau et al., 2008; Daste et al., 2015; Gupton and Gertler, 2010; Han et al., 2017; Jahn and Scheller, 2006; Martinez-Arca et al., 2001b; Martinez-Arca et al., 2003; Proux-Gillardeaux et al., 2005; Proux-Gillardeaux et al., 2007; Vats and Galli, 2022; Verderio et al., 2012; Wang et al., 2018) (Fig. 2A). To account for different image qualities and/or fusion reporter signal intensity (Fig. 2A), we iteratively refined the detection threshold values kσ_wavelet_ (Fig. 1B and Table 2), dF/σ_dF_ (Fig. 1D and Table 4) and the time window centered around candidate events at t_0_ (Expanding frames; Fig. 1D and Table 4) until we reach a plateau of detected TSPAN- and v-SNARE-reported fusion events. While detection thresholds allow for distinct fusion intensity with respect to the local background, our algorithm explores a user-set time window to optimize the capture of fusion events of different dynamics. No further adjustments were made before running the identification process of each vesicle population in a batch-processed manner. We eventually reported a total number of 481 and 315 analyzed events for TSPAN and v-SNARE population respectively. After in-depth reviews, we removed 10 TSPAN-pHluorin and 2 v-SNARE-pHluorin detection hits that did not correspond to fusion events but rather to filopodia tips coming in and out of focus, extracellular fluorescent objects and stationary vesicles. The overall identification error rate of 1.5% is comparable or better than previously published algorithms (Urbina et al., 2018; Yuan et al., 2015).

**Figure 2.**
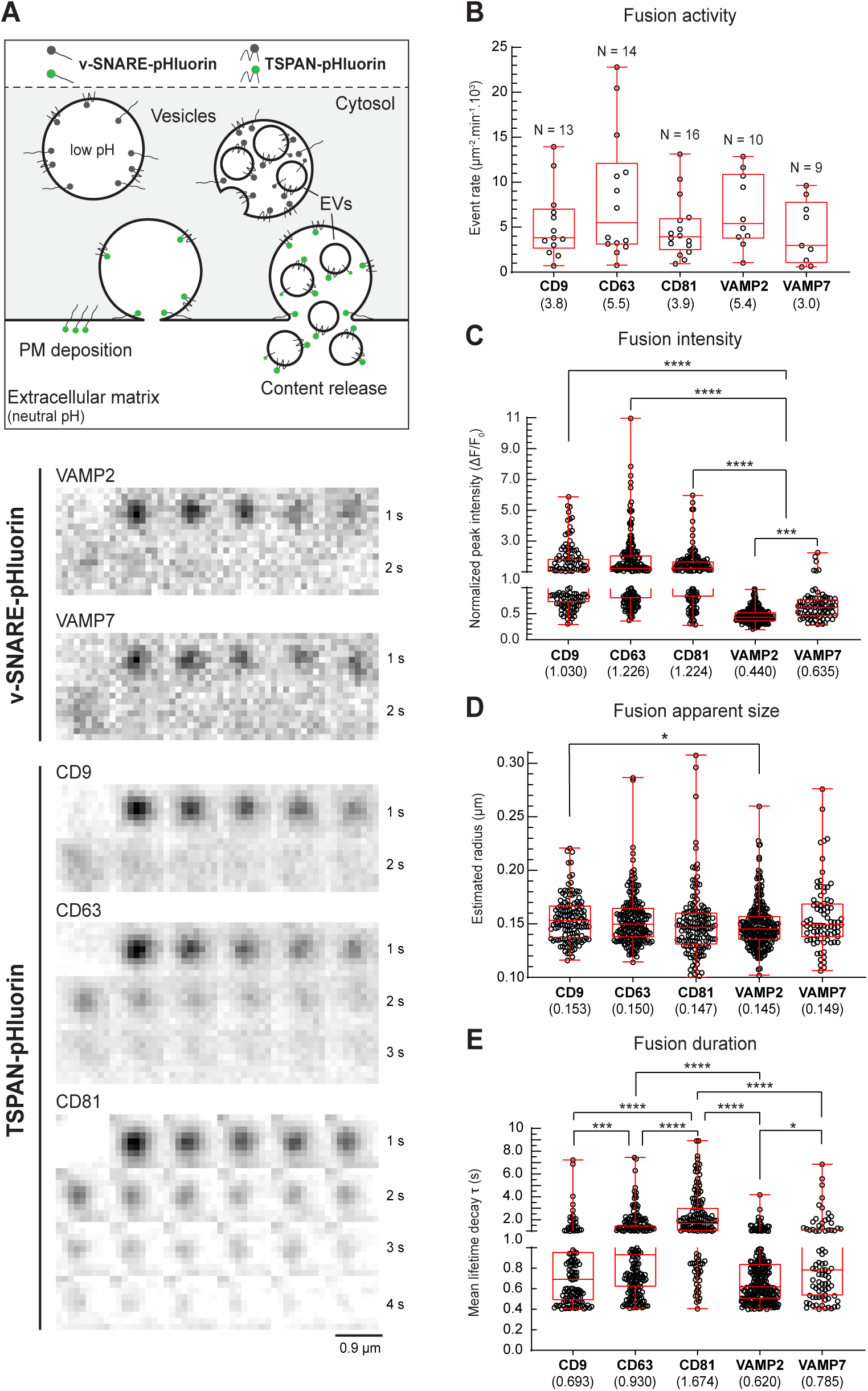
Feature evaluation of fusion events evidences distinct dynamics of vesicle-mediated exocytosis. **(A)** Schematic model illustrating the markers used in this work to monitor vesicle fusion to the PM. The content and/or the vesicle membrane are labeled with pHluorin. Time-lapse montages of TSPAN- and v-SNARE-mediated fusion events are shown (invert LUT). **(B-E)** Combined scatter dot (black circle) and box and whisker (red border) plots are shown for individual population of labeled vesicles. The median values of fusion **(B)** activity, **(C)** normalized peak intensity (ΔF/F_0_), **(D)** apparent size and **(E)** event duration (τ) are displayed in brackets. **(B)** Number of analyzed cells (N) for TSPAN and v-SNARE proteins are indicated and represented with black circles. **(C-E)** Feature comparisons between TSPAN- and v-SNARE-labeled vesicular population were performed using a Kruskal-wallis’s test followed with a post hoc Dunn’s multiple comparison test. Only statistical differences are indicated. Black circles correspond to single fusion events. **(C)** For fusion activity reported by ΔF/F_0_: ****, P < 0.0001; ***, P < 0.0002; **(D)** For fusion apparent size, *, P < 0.04; **(E)** For fusion duration reported by the mean lifetime τ, ****, P < 0.0001; ***, P < 0.0002; *, P < 0.03.

The capacity to modulate the identification algorithm ensures our plugin to be adaptive in detecting distinct types of fluorescently-reported fusion event with high accuracy.

### ExoJ feature evaluation provides quantitative insights into distinct forms of vesicle-mediated exocytosis

We next evaluated whether ExoJ could be used to quantitatively study exocytosis. TSPAN CD9, CD63 and CD81 are known to form protein microdomains at the PM (Hemler, 2005; le Naour et al., 2006) regulating various cellular function ranging from cell adhesion to proliferation (Becic et al., 2022) and signaling at the immunological synapse (Levy and Shoham, 2005). Recently, these three TSPANs are found to be enriched to various degrees in the membrane of EV subpopulations such as ectosomes and exosomes (Crescitelli et al., 2013; Escola et al., 1998; Kowal et al., 2016; Mathieu et al., 2021; Théry et al., 2018; Verweij et al., 2018). Exosomes are formed as intraluminal vesicles in late endosomes, and their secretion can be detected using pHluorin-tagged proteins (Fig. 2A). Previous studies have, however, found CD9 mainly enriched in other subtypes of EVs directly budding from the plasma membrane and hence not monitored by our approach (Kowal et al., 2016; Mathieu et al., 2021; Verweij et al., 2018). We examined post-fusion features of TSPAN subpopulations and compared them to transport vesicles reported with v-SNARE-pHluorin (Fig. 2A). In Hela cells, the detection algorithm recorded a statistically similar fusion activity between individual reporter populations (Fig. 2B) with no correlation with the intensity and fusion apparent size (Fig. 2C,D). Despite TSPAN-pHluorin-reported events displaying two to three-fold higher fusion intensity than the v-SNAREs, our unbiased automatic detection method was able to capture exocytic events reported by both, highlighting the effectiveness of our approach even in the presence of significantly varying signal intensity (Fig. 2A,C). The recorded difference among v-SNARE-pHluorin subpopulations is consistent with previous findings in various cellular contexts, where v-SNAREs VAMP2 and VAMP7 were observed to be differently localized in endosomes at different stages (Gupton and Gertler, 2010; Wang et al., 2018) (Fig. 2C,E). The estimation of apparent size of events reported by TSPAN-pHluorin (Fig. 2D) matches previous qualitative analysis at supra-optical electron microscopy (EM) resolution using a dynamic correlative light electron microscopy approach (Verweij et al., 2018). In addition, the apparent size did not differ significantly from v-SNARE-pHluorin-reported events (Fig. 2D) which was at the lower end of the total spectrum previously reported by Altick and colleagues (Altick et al., 2009). This, however, closely matches early quantitative EM studies in the central nervous system (Roizin et al., 1967). Focusing on the fusion duration, we first noted the short-lived fluorescence signal of v-SNARE-pHluorin, corresponding to a rapid either post-fusion lateral diffusion at the PM or endocytic process (Alberts et al., 2006; Miesenböck et al., 1998; Urbina et al., 2018; Verweij et al., 2018; Wang et al., 2018) (Fig. 2E). We also showed that subpopulations of TSPAN-pHluorin displayed a significantly distinct fusion event duration (Fig. 2E). The signal duration at the PM of CD81-pHluorin was almost two- and three-fold longer than CD63- and CD9-pHluorin respectively. This could not previously be measured with this level of accuracy (Verweij et al., 2018). The divergence in fusion duration could not be explained by a noticeable difference in apparent size, nor a significantly higher amount of TSPANs (Fig. 2C,D). For CD63 and CD81, we hypothesized that this difference could either reflect the various regulatory fusion machineries and/or the heterogeneity in the composition, the biogenesis, the maturation and the secretion of reported EV subpopulations (Edgar et al., 2016; Larios et al., 2020). Indeed, TSPANs are known to organize in distinct TSPAN-enriched microdomains that consist primarily of less than 120 nm homotypic TSPAN-TSPAN interactions and specific partner proteins (Charrin et al., 2009; Rubinstein et al., 1996; van Deventer et al., 2021) and mark distinct populations of exosomes (Matsui et al., 2021). Furthermore, the signal duration of CD9-pHluorin was similar to v-SNARE-labeled population of vesicles, as previously observed (Verweij et al., 2018). This suggested that CD9-pHluorin bursts of fluorescence may reflect post-fusion lateral diffusion over the PM or endocytosis similarly to v-SNARE-pHluorin rather than exosome secretion in most events.

Altogether, our computer vision-assisted tool enabled us to record and evaluate features of different types of cargo and/or vesicles, providing quantitative insights into post-fusion dynamics.

### Performance assessment of ExoJ evidences a highly robust solution to detect fusion events

To fully assess the performance of ExoJ, we simulated movies with synthetic data corresponding to fluorescence signal of fusion events randomly distributed at the cell surface (Fig. 3A). In our simulation, we modeled a random number of events across a wide range of normalized intensity ΔF/F_0_, apparent size and duration τ features including data collected in this study and in different cellular contexts and/or protein reporters from others (Persoon et al., 2019; Urbina et al., 2018; Verweij et al., 2018; Wang et al., 2018; Yuan et al., 2015). We also simulated events featured as non-relevant, exhibiting distinct signal decay behaviors (see Materials and Methods for details). We additionally tested the influence of increasing Gaussian noise signal which coarsely recapitulates local background variation (Fig. 3A). ExoJ was able to accurately capture simulated events within the range of typical signal-to-noise ratio observed experimentally, with an identification error rate (also equates to 100 – accuracy) as low as 1.5% up to 3% which remains comparable to Urbina and colleagues’ work (Urbina et al., 2018) (Fig. 3B). At each noise increment, we thoroughly adjusted the algorithm requirements by refining kσ_wavelet_ (step 1), dF/σ_dF_ (step 3), both goodness-of-fit R^2^ and the time window to optimize event identification (Fig. 1D and Table 4). In response to incremental Gaussian noise, ExoJ detection capability was significantly degraded with an error rate up to 7% ± 1.4%, which still performs at a higher level of accuracy under equal noise condition (Urbina et al., 2018). Error rate alone is an incomplete measure on simulated movies with disparate fusion activity. Thus, we introduced standard metrics such as sensitivity, precision and specificity which altogether score the ability to correctly detect events while accurately discarding those featured as non-relevant (see Materials and Methods for details). Results on simulated events demonstrated that ExoJ is a highly robust tool with scoring metrics above 97% (Fig. 3C) which bettered previously published algorithms (Sebastian et al., 2006; Urbina et al., 2018). Increasing the contribution of Gaussian noise signal significantly impaired the ability to correctly capture fusion events down to 89.6% (sensitivity) in the lowest condition. However, these captured events were accurately identified (precision of 97.8%) while avoiding those non-relevant (specificity of 97.3%) (Fig. 3C). This balance between sensitivity and precision was observed independently of added Gaussian noise signal, and reported with a very high F1 score (Fig. 3C). In addition, there was no correlation between ExoJ scoring results and the frequency rate of fusion events simulated per movie. The flexibility of ExoJ enabled us to relax the algorithm requirements to account for changes in imaging signal-to-noise ratio without degrading its ability to effectively capture fusion events.

**Figure 3.**
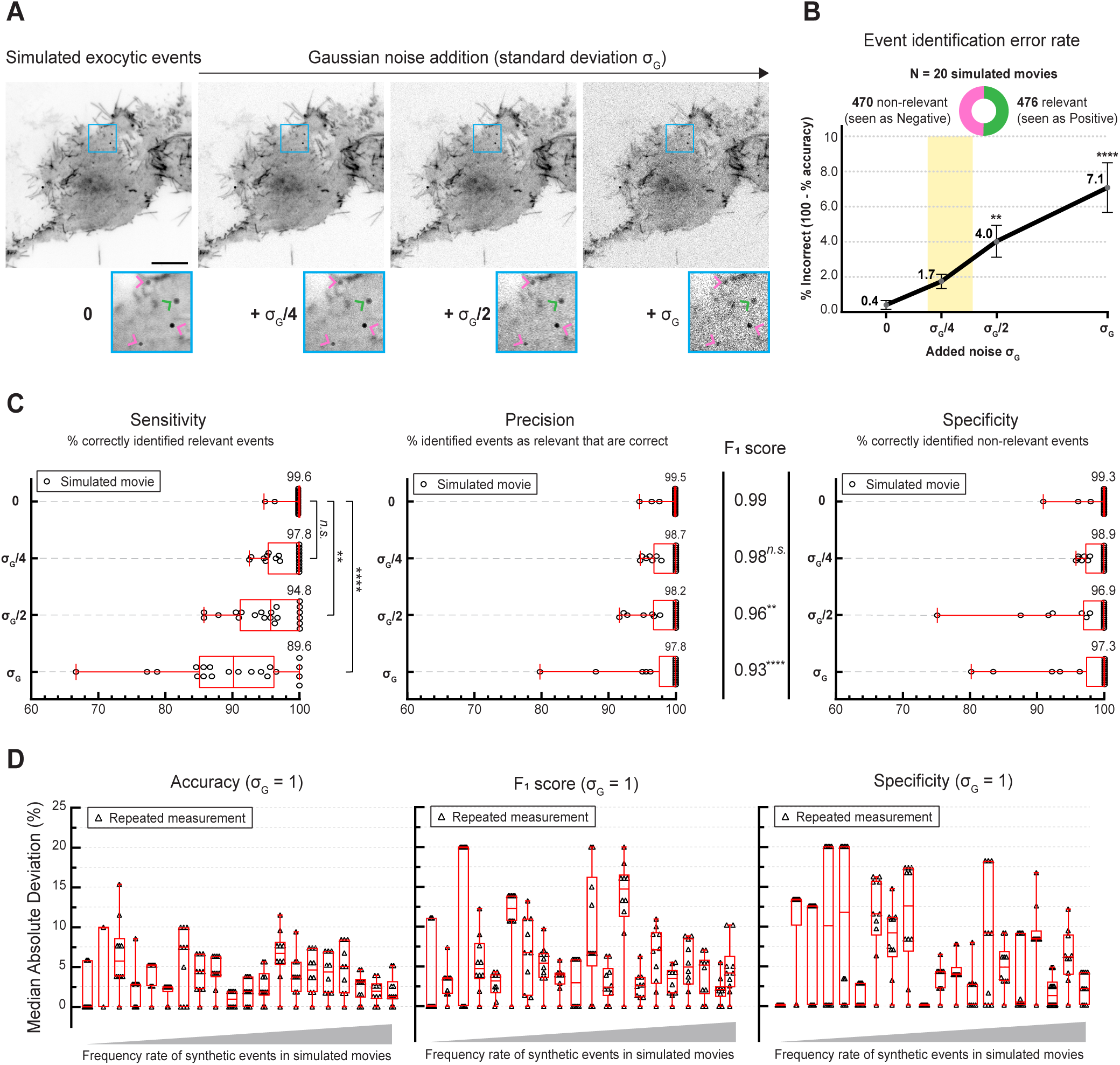
ExoJ is a highly robust tool to identify distinct forms of fusion events. **(A)** Examples of simulated exocytic event signal at the cell surface with increasing Gaussian noise intensity (Invert LUT). The inset (blue box) shows four simulated relevant (resp. non-relevant) events indicated by green (resp. pink) arrowhead(s). The total number of simulated exocytic events as well as their XY location, normalized peak intensity ΔF/F_0_, apparent size and duration τ are randomly set (see Materials and Methods). **(B)** A total of 20 movies were simulated with an overall balanced number of randomly generated relevant (green) and non-relevant exocytic (pink) events as indicated. The XY plot corresponds to the evaluation of identification error rate with increasing Gaussian noise intensity. The estimated noise range of experimental image series is indicated in yellow. Data represent mean ± SEM. Kruskal-wallis’s test was performed (P < 0.0001) followed with a post hoc Dunn’s multiple comparison test with 0 as the control (****, P < 0.0001; **, P < 0.002; n.s., non-significant). **(C)** Assessment of ExoJ sensitivity (proportion of correctly identified relevant events), precision (proportion of identified events as relevant ones that were actually correct), specificity (proportion of correctly identified non-relevant events) with increasing Gaussian noise intensity are shown as combined scatter dot (black) and box and whisker (red) plots. Mean value as well as average F_1_ score are reported for each condition. Black dots correspond to the scoring results of individual simulated movies. Kruskal-wallis’s test was performed (P < 0.0001) followed with a post hoc Dunn’s multiple comparison test with 0 as the control (****, P < 0.0001; **, P < 0.002; n.s., non-significant). **(D)** Assessment of ExoJ repeatability on individual simulated movies with added Gaussian noise of sigma equal to 1, and reported by the MAD of accuracy, F_1_ score and specificity. Scoring results of simulated results are shown as combined scatter dot (black) and whisker (red) plots, and were sorted according to the simulated fusion activity.

Finally, we asked whether the event detection procedure is reliable enough to handle biological variability between individual cells. To mimic this, we took advantage of the random spatial distribution of Gaussian noise added to simulated timeseries. We added noise with a sigma value of 1, followed by event identification using constant set of parameters previously optimized for single simulated movies. We repeated this procedure ten times, scored ExoJ performance at each round and reported the overall relative variation using the MAD. Our results demonstrated a high repeatability in detection, independently of the simulated fusion event rate as well (Fig. 3D). The average MAD evaluation showed an overall variation of 4.1%, 4.0% and 7.7% for accuracy, F_1_ score, and specificity, respectively.

Taken together, assessment of ExoJ performance revealed a robust tool in identifying fusion events insensitive to experimental noise.

## Discussion

Here we presented ExoJ, a computer-vision assisted tool for the automated identification and analysis of vesicle exocytosis events marked by a pH-sensitive probe. The identification algorithm was designed as a fully user-configurable processing pipeline, making it suitable to detect and analyze various forms of exocytic events independently of the imaging condition. Built-in options were implemented for visualization and manual curation of event features.

### Considerations and future applications

We demonstrated ExoJ ability in monitoring fusion events reported by TSPAN- and v-SNARE-pHluorin proteins with high accuracy. Reporting of spatiotemporal features provided quantitative insights with unique details between and among subpopulations of vesicle exocytosis. Using simulated data covering a wide range of feature data, performance assessment underscored ExoJ as a robust and reliable identification tool. As it is, ExoJ usage has no specific constraints on vesicle exocytosis features including normalized intensity ΔF/F_0_, duration of signal decay τ and apparent size. In response to changing condition in imaging, the sensitivity of event detection could be adjusted by lowering the detection threshold value kσ right from the first step (Fig. 1B). This will increase the number of captured spots along with the processing time in the subsequent steps (up to 12 minutes per movie on average for +σ_G_ condition; Fig. 3A). We derived the duration of fusion events by evaluating the mean lifetime τ of fluorescence peak intensity profiles F fitted with single mono-exponential decay functions (Fig. 1A). While most studies described exocytosis kinetics with single exponential decay, raw data are available for export and manual curation using a different mathematical model as performed by Mahmood and colleagues (Mahmood et al., 2023). Apparent size is not a limiting factor since spots are seen as gaussian-shaped intensity over a user-defined range of gaussian widths. In contrast to fusion intensity and duration, apparent size results were, however, modest due to the resolution limit of TIRFM (Fig. 2D). The advent of fast-imaging Structured Illumination Microscopy and novel fluorescent markers may improve the recording of the structural dynamics of exocytosis with unprecedented resolution (Huang et al., 2018; Li et al., 2015; Liu et al., 2021b; Roth et al., 2020). In combination with ExoJ, we are confident that these developments will certainly help providing insights into exocytosis-associated protein dynamics, and exploring machinery involved in EV biogenesis and secretion (Verweij et al., 2022).

### Relevance to existing methods

Numerous bioinformatics tools on single or cross-platforms have previously been developed for specific applications which make fair comparison difficult (Bebelman et al., 2020; Burchfield et al., 2010; Diaz et al., 2010; Jullie et al., 2014; Mahmood et al., 2023; Moro et al., 2021a; Persoon et al., 2019; Sebastian et al., 2006; Urbina et al., 2018; Wang et al., 2018; Yuan et al., 2015). In addition, some image formats are not accepted, file size limit could hinder the time of analysis, and feature extraction approach could vary (Mahmood et al., 2023; Urbina and Gupton, 2021; Urbina et al., 2018; Yuan et al., 2015). Our goal in developing ExoJ was to provide a common, yet robust computer-vision assisted solution regardless of the experimental condition. To this end, we designed a GUI-based tool as a plugin for ImageJ2/Fiji, a well-established platform for biological image analysis (Schindelin et al., 2012). As a result, ExoJ accepts all type of image formats and bit depths recognized by ImageJ2/Fiji. Prior to running ExoJ if necessary, experimental data could be pre-processed using built-in filter toolboxes. In the last decade, supervised methods approaches have achieved great success in providing solutions for detecting subcellular structures in fluorescent microscopy images (Boland and Murphy, 2001; Hu et al., 2010; Johnson et al., 2015). In particular, machine learning approaches were successfully applied for spot detection upon a training phase (Jiang et al., 2007; Lin et al., 2019). Careful selection and label of training datasets is, however, an essential prerequisite, either manually or using other detection methods. Instead, we opted for unsupervised methods which also perform well with simple user-configurable features such as fluorescently patterned vesicular spots (Basset et al., 2015; Smal et al., 2010). We illustrated the capacity of ExoJ to detect distinct exocytic events, illustrated here by TSPAN- and SNARE-pHluorin, using a limited number of biological parameters (Fig. 1B-D). We based ExoJ detection on a proven wavelet transform algorithm (Lagache et al., 2018; Olivo-Marin, 2002; Püspöki et al., 2016; Ruusuvuori et al., 2010; Toonen et al., 2006; Yuan et al., 2015). We also introduced the statistical measure MAD to hone the capture of candidate exocytic events which could partly explain the performance difference with Yuan and colleagues’ wavelet-based tool (Yuan et al., 2015). Our detection algorithm also explored a user-set time window centered around exocytic events, whereas previously published tools had it fixed or hardly modifiable (Mahmood et al., 2023; Urbina and Gupton, 2021; Urbina et al., 2018; Yuan et al., 2015). Throughout the workflow, users can go back and forth to readjust parameters and preview results. Much efforts have been devoted to facilitate the identification of user-defined genuine events as well as implementing options for data visualization, manual curation and export (Fig. S1). In addition to our study, ExoJ has recently demonstrated its capability in providing quantitative insights on the role of SNARE protein SNAP29 in CD63-pHluorin-labelled EVs in PC-3 cells imaged with a confocal spinning-disk microscope (Hessvik et al., 2023). We are convinced that ExoJ could become a standard tool for quantitative review of comparative studies of vesicle exocytosis in an unbiased manner.

## Materials and Methods

### Installation and system requirements

ExoJ was developed as an ImageJ2/Fiji (Schindelin et al., 2012) plugin for the automated detection and analysis of exocytic events in 8 or 16-bit grayscale fluorescent image series. The plugin is published under the GPLv3 license. It uses ImageJ2/Fiji capabilities to open a wide range of image formats using the plugin Bio-Formats (Linkert et al., 2010). The plugin ExoJ can be downloaded on the following website https://www.project-exoj.com/ with the code source available at https://github.com/zs6e/excytosis-analyzer-plugin. To install the plugin, place the .jar file in the ImageJ2/Fiji plugin directory. After a restart, ExoJ will be available in the ImageJ2/Fiji plugin menu. ExoJ is compatible with ImageJ v1.53t or newer version, and has been tested on Windows and MacOS platforms. We recommend using a 64-bit operating system with Java 8 installed and at least 4 GB RAM.

### Photobleaching correction

The main challenges posed by reliable vesicle detection are associated with prolonged live-cell acquisition, such as cell migration, focus drift and photobleaching. While cell migration and focus drift lead to unpredictable fluorescent changes at a given pixel, the photobleaching effect can be theoretically compensated to a certain extent (Miura, 2020). The variations of fluorescent signals over time within cell compartments are occurring at slower dynamics compared to single fusion events. To compensate for this effect, we implement an optional custom-written correction based on a non-fitting, pixel-by-pixel weighted detrending method. The option is available during the spot detection step (Fig. 1B). If “Correct photobleaching” is selected in the setting menu, the correction is applied on the whole time series and can be reversed upon deselection. In detail, for each pixel P_xy_, we measured the fluorescence intensity difference between the first P_xy0_ and last P_xyt_ frame in the image series. These values were then normalized by the difference between the maximal μ_max_ and minimal μ_min_ intensity, also calculated throughout the image series to generate a weight map (W_xy_) such as:

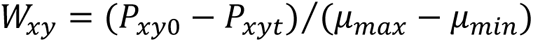

Next, for each frame i we calculate the difference d_i_ between the mean intensity of the entire image series μ and the mean intensity of that frame μ_i_. Finally, the fluorescence intensity of every pixel at each frame is compensated by W_xy_ d_i_ to obtain the final detrended image series.

### Built-in tools

A series of built-in options are implemented to visualize single and/or the population of fusing vesicles. Each analyzed fusion event can be visualized as a sequence of cropped images with associated fluorescence peak intensity F, 1^st^ order differential intensity dF and estimated radius fitting plots. We also record the fluorescence spreading over time associated with each vesicle fusion event (Spatial dynamics; Fig. S1A) as initially proposed by Bowser and colleagues (Bowser and Khakh, 2007). We allow the possibility to discard events using the appropriate button in the result table (Remove; Fig. S1A). Similarly, candidate fusion events can be manually added and immediately reviewed (draw and press Add; Fig. S1A). Upon selection, the plugin will prompt the user to define the timeframe of candidate fusion events. A spatiotemporal map distribution of fusion events is available upon selection in the result table (Fig. S1B), and can further be exported as an RGB image.

### Reporters of fusion events in live cells

We used TSPAN and v-SNARE proteins as fusion event reporters. Specifically, we focused on TSPAN CD9, CD63 and CD81 (Escola et al., 1998; Kowal et al., 2016; Théry et al., 2018; Yáñez-Mó et al., 2015), and v-SNARE VAMP2/synaptobrevin2 (vesicle-associated membrane protein 2) and VAMP7/TI-VAMP (vesicle-associated membrane protein 7 / Tetanus neurotoxin-Insensitive vesicle-associated membrane protein) (Chaineau et al., 2009; Galli et al., 1998) proteins coupled to pHluorin (Alberts et al., 2006; Miesenböck et al., 1998). CD9-pHluorin, CD63-pHluorin and CD81-pHluorin plasmids were constructed as previously described by Verweij and colleagues (Verweij et al., 2018). VAMP2-pHluorin was a kind gift from Dr. T. Ryan (Cornell University, USA). VAMP7-pHluorin construction corresponds to an improved version (Chaineau et al., 2008) previously described (Alberts et al., 2006; Martinez-Arca et al., 2000) and further characterized (Burgo et al., 2013; Wang et al., 2018). We used Hela cells cultured in DMEM supplemented with 10% FBS (Perbio Sciences; HyClone), 100 U/ml penicillin G, 100 mg/ml streptomycin sulfate and 2 mM glutamax (Invitrogen, Thermofischer Scientific). Cells at 50% to 60% confluence were transfected using Lipofectamine 2000 reagent (Invitrogen) on either 35-mm glass bottom petri dishes (Ibidi) scaled with 500 ng of TSPAN-plasmids or 18-mm round glass coverslips deposited in 12-well plates scaled with 1 μg of VAMP plasmid. Glass coverslips were further mounted on a Chamlide EC magnetic chamber (LCI).

### Live cell imaging

Prior to imaging, cell medium was replaced with homemade Hepes-buffered Krebs-Ringer as previously used by Danglot and colleagues (Danglot et al., 2012) or Leibovitz’s L-15 solution (Gibco). Hela cells were imaged 24 h after transfection on an inverted microscope (Axio Observer 7, Zeiss) equipped with a TIRF module and a top stage imaging chamber (Tokai Hit STX-CO2) ensuring a constant temperature at 37°C. All imaging experiments were carried out with a 63x 1.4 NA oil objective (Zeiss), an air-cooled 488 nm laser line at 0.4-0.6% power, with a TIRF angle set for 300-400-nm penetration depth and an additional optovar 1.6x to reach a pixel size of 100 nm. Images were acquired with Zen Black (SP2.3, Zeiss) onto an electron-multiplying charge-coupled device camera (iXon, Andor) at a frame rate of 5 Hz. Fluorescent timeseries were saved as tiff or czi files. Fusion activity was defined as the number of detected events throughout the cell surface over the course of a time-lapse experiment, which was typically 3 minutes (Fig. 2B). Thus, experimental data in this study consisted in single fluorescent timeseries of 901 images. Cell surface measurement was carried out using ImageJ/Fiji segmentation tools (Schindelin et al., 2012). Number of analyzed cells (N) per labeled population is indicated.

### Procedure for TSPAN- and vSNARE-reported fusion events

The detection parameters are adapted to the different types of fusion event. After a few rounds of optimization, we set the detection threshold values kσ_wavelet_ to 4.0-5.0 (resp. 3.0) and dF/σ_dF_ to 4.0-5.0 (resp. 3.0-3.5) for fusion events reported by TSPAN-pHluorin (resp. v-SNARE-pHluorin). We imposed a minimal signal duration of two frames (Fig. 1D). We completed the detection of candidate events with additional frames corresponding to 1 s (resp. 3-4 s) before (resp. after) the maximal peak fluorescence at t_0_ (Expanding frames; Fig. 1C and Table 4). For v-SNARE-pHluorin, we limited the temporal window to 0.6 s (resp. 2-3 s) before (resp. after) the maximal peak. The fluorescence profiles were fitted with a minimal number of five points and a minimal goodness-of-fit R^2^ of 0.75 were set as the fitting threshold. The initial vesicle radius (step 1; Fig. 1B) and the tracking parameters (step 2; Fig. 1C) do not vary between TSPAN and v-SNARE protein reporters. In details, we reported the number of identified events n for each protein reporters as follows: CD9, n = 129; CD63, n = 200; CD81, n = 152; Vamp2, n = 232; Vamp7, n = 83.

### Computer simulation of exocytosis

To fairly assess the robustness of ExoJ, we generated 20 movies of 901 images with randomly-distributed gaussian-shaped pHluorin signal intensity on the cell surface. We simulated two groups of synthetic events, labeled as relevant and non-relevant according to the features we imposed. A series of parameters was defined to encompass distinct features of relevant exocytic events including those approximated by users in this study as well as previous works (Persoon et al., 2019; Urbina et al., 2018; Verweij et al., 2018; Wang et al., 2018; Yuan et al., 2015). In details, we imposed to the simulated events a random distribution of: (1) normalized fluorescence peak intensity ΔF/F_0_ as low as 10% up to 200% above the local background (centered circle of 9-pixel radius) and (2) apparent size over a range of gaussian widths from 3 to 7 pixels. Events featured as relevant exhibited single exponential decay behaviour with a mean lifetime τ from 1 to 50 timeframes. In contrast, we modified the transient nature of exocytic events featured as non-relevant where the signal decay could be (1) 1-frame long, (2) long-lived and constant fluorescence peak intensity, or (3) a damped sine wave along with (4) a random spatial displacement. The maximal number of simulated relevant and non-relevant events was both fixed to 50, and randomly set for each movie. The simulated fusion activity eventually ranged from 1.2 to 9.8 μm^-2^.min^-2^.10^3^ which matches with previously recorded frequency rate (Persoon et al., 2019; Urbina et al., 2018; Verweij et al., 2018; Wang et al., 2018; Yuan et al., 2015). The identification pipeline was optimized to capture simulated events labeled as relevant. To represent random noise, we added a range of fluorescent intensities that follow a Gaussian distribution to the simulated movies using ImageJ built-in function Add Specified Noise.

### Performance evaluation metrics

To benchmark the performance of ExoJ several metrics were used including identification error rate, sensitivity, precision, specificity and F1 score. These metrics were recorded for analysis on movies with simulated events with relevant (resp. non-relevant) features considered as true positive (TP) (resp. true negative, TN) attribute. When running ExoJ, each identified event was considered with either relevant (TP) or non-relevant (labeled as false positive, FP) attributes. Finally, undetected relevant event was considered with false negative (FN) attribute. Considering these definitions, metrics are measured based on direct count of true and false positives and negatives such as:

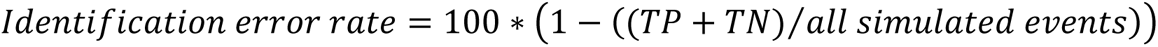

which corresponds to the proportion of correctly identified relevant and non-relevant events out of all the simulated events.

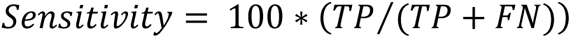

which reports the proportion of correctly identified relevant events out of the total number of simulated relevant events;

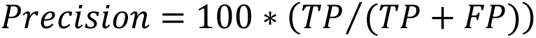

which reports the proportion of correctly identified relevant events out of the total number of identified fusion events;

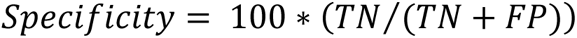

which reports the proportion of correctly identified non-relevant events out of the total number of simulated non-relevant events.

We additionally introduced F_1_ score (Fawcett, 2006) which reports the capability to both capture relevant events (sensitivity) and be accurate with the events ExoJ does capture (precision) as:

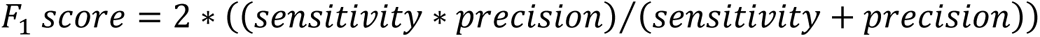

with 1.0 as the highest possible value.

### Statistical data analysis

We performed statistical analysis using Prism 9.0 (GraphPad). For multiple comparisons between populations of labeled vesicles, Kruskal-Wallis non-parametric ANOVA was used to determine significance, followed by Dunn’s post hoc test for multiple comparisons between populations of labeled vesicles and simulated movies with increasing Gaussian noise intensity. Statistical significance was considered for α = 0.05. Data are reported as box and whisker plots. Boxes show the median +/-interquartile range and the whiskers go down to the minimum and up to the maximal value in each dataset. For each population, median value is displayed in brackets below each corresponding box and whisker plot unless mentioned otherwise in the legend.

## Acknowledgements

We acknowledge the NeurImag imaging facility of the Institute of Psychiatry and Neurosciences of Paris (Inserm U1266) where the imaging experiments were carried out. NeurImag is part of GIS IBISA and the national infrastructure France BioImaging supported by the French National Research Agency (ANR-10-INBS-04). We thank Dr. Liangyi Chen and Dr. Yongdeng Zhang for sharing their Matlab executable tool for exocytosis detection. We are grateful to Julie N’guyen for maintaining and providing HeLa cells. We thank Dr. Sebastien Nola, Dr. Cédric Blouin and Brett Davis for their insightful comments on the manuscript. Zeiss Confocal LSM 880 Elyra PS.1 equipped with a TIRF module was purchased with a Région Ile-de-France grant awarded to Lydia Danglot DIM Cerveau et Pensée project. Guillaume Van Niel would like to acknowledge Institut National du Cancer, Fondation ARC pour la Recherche sur le Cancer and the ANR for their financial support.

## Author contributions

**JL**: Conceptualization, Methodology, Software, Data curation, Resources, Visualization, Writing-Original draft, Writing-Review and Editing. **FJV**: Methodology, Resources, Investigation, Validation, Writing-Review and Editing. **GVN**: Supervision, Funding acquisition, Writing-Review and Editing. **TG**: Resources, Writing-Review and Editing. **LD**: Conceptualization, Supervision, Methodology, Funding acquisition, Resources, Investigation, Writing-Review and Editing. **PB**: Conceptualization, Supervision, Project administration, Methodology, Software, Data curation, Investigation, Formal analysis, Visualization, Writing-Original draft, Writing-Review and Editing.

## Availability of data and Material

The datasets used and/or analysed during the current study are available from the corresponding authors on reasonable request. Time series including the one used in this report as well as five simulated movies are available in the Zenodo repository: https://doi.org/10.5281/zenodo.6610894 and https://doi.org/10.5281/zenodo.7595198. The latest version of ExoJ as well as a tutorial can be found on the following website https://www.project-exoj.com/. The source code was deposited in a GitHub repository: https://github.com/zs6e/excytosis-analyzer-plugin.

## Abbreviations

PM: Plasma membrane
TIRF: Total Internal Reflection Fluorescence
FP: Fluorescent protein
MAD: Median absolute deviation
RGB: Red green blue
VAMP2: Vesicle-associated membrane protein 2
VAMP7/TI-VAMP: Vesicle-associated membrane protein 7 / Tetanus neurotoxin-Insensitive vesicle-associated membrane protein
EV: Extracellular vesicle
EM: Electron microscopy
SIM: Structured illumination microscopy

---

**Figure S1.**
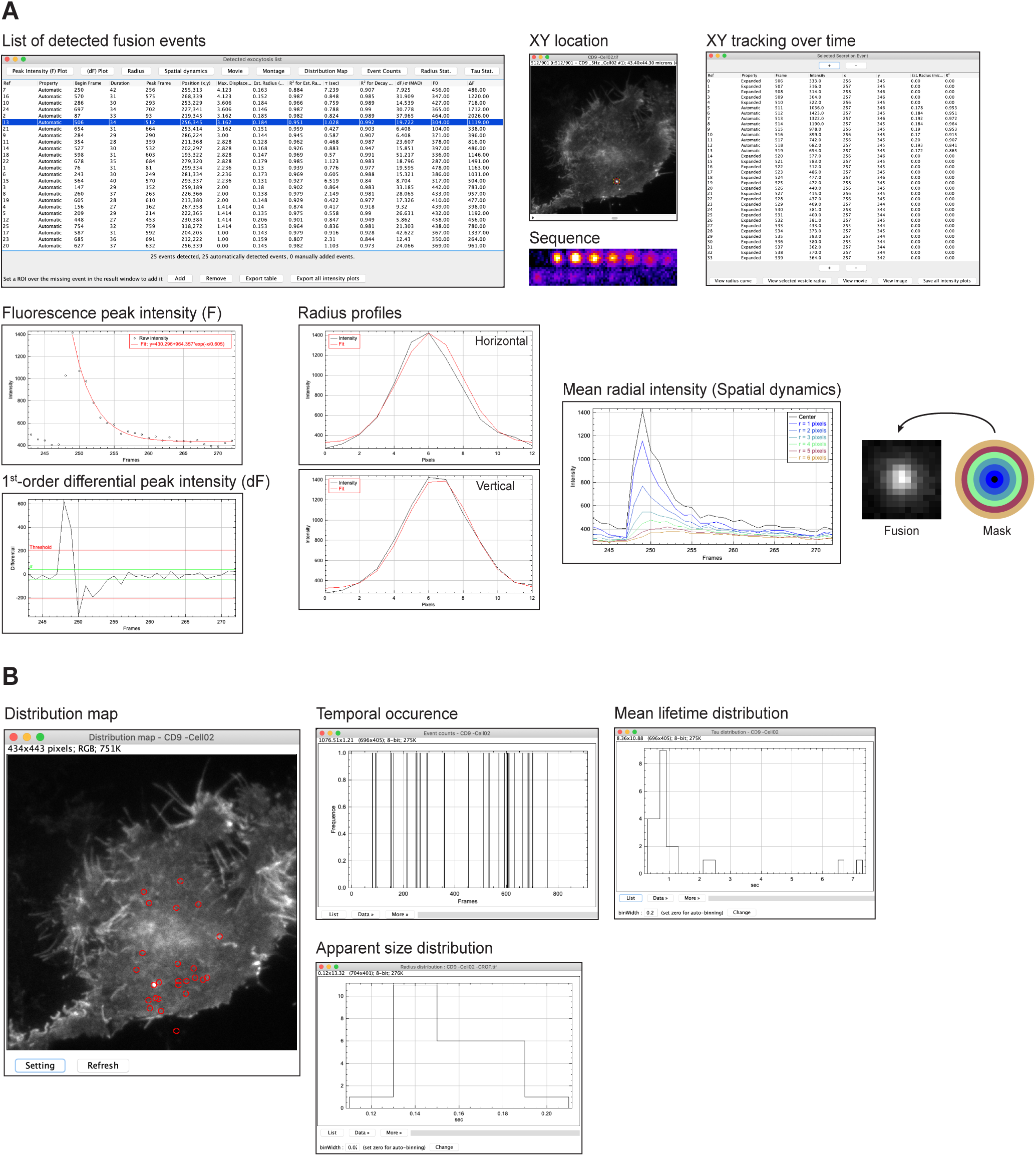
Built-in tools for visualizing fusion events. **(A)** Series of measurements on single fusion events are summarized in a result table, including the XY location, the onset time (peak frame), the maximal displacement during the fusion event, the detection threshold value (dF/σ_dF_), the local background (F_0_) and normalized peak fluorescence intensity (ΔF/F_0_), the estimated apparent size (est. radius) and mean lifetime decay (τ). Corresponding graphs can be displayed for each fusion event upon selection in the result table. **(B)** Visualization of the distribution of fusion event duration (Tau Stat.), apparent size (Radius Stat.) and temporal occurrence (Event counts) throughout the fluorescent times series can be generated upon selection. A spatial occurrence map (Distribution map) is also available for export.

## References

Alberts, P., Rudge, R., Irinopoulou, T., Danglot, L., Gauthier-Rouvière, C. and Galli, T. (2006). Cdc42 and Actin Control Polarized Expression of TI-VAMP Vesicles to Neuronal Growth Cones and Their Fusion with the Plasma Membrane. Mol Biol Cell 17, 1194–1203.

Altick, A. L., Baryshnikova, L. M., Vu, T. Q. and von Bartheld, C. S. (2009). Quantitative analysis of multivesicular bodies (MVBs) in the hypoglossal nerve: Evidence that neurotrophic factors do not use MVBs for retrograde axonal transport. J Comp Neurol 514, 641–657.

Axelrod, D. and Omann, G. M. (2006). Combinatorial microscopy. Nat Rev Mol Cell Biol 7, 944–952.

Basset, A., Boulanger, J., Salamero, J., Bouthemy, P. and Kervrann, C. (2015). Adaptive Spot Detection With Optimal Scale Selection in Fluorescence Microscopy Images. IEEE Transactions on Image Processing 24, 4512–4527.

Bebelman, M. P., Smit, M. J., Pegtel, D. M. and Baglio, S. R. (2018). Biogenesis and function of extracellular vesicles in cancer. Pharmacol Ther 188, 1–11.

Bebelman, M. P., Bun, P., Huveneers, S., van Niel, G., Pegtel, D. M. and Verweij, F. J. (2020). Real-time imaging of multivesicular body–plasma membrane fusion to quantify exosome release from single cells. Nat Protoc 15, 102–121.

Becic, A., Leifeld, J., Shaukat, J. and Hollmann, M. (2022). Tetraspanins as Potential Modulators of Glutamatergic Synaptic Function. Front Mol Neurosci 14,.

Boland, M. V. and Murphy, R. F. (2001). A neural network classifier capable of recognizing the patterns of all major subcellular structures in fluorescence microscope images of HeLa cells. Bioinformatics 17, 1213–1223.

Bowser, D. N. and Khakh, B. S. (2007). Two forms of single-vesicle astrocyte exocytosis imaged with total internal reflection fluorescence microscopy. Proceedings of the National Academy of Sciences 104, 4212–4217.

Burchfield, J. G., Lopez, J. A., Mele, K., Vallotton, P. and Hughes, W. E. (2010). Exocytotic Vesicle Behaviour Assessed by Total Internal Reflection Fluorescence Microscopy. Traffic 11, 429–439.

Burgo, A., Proux-Gillardeaux, V., Sotirakis, E., Bun, P., Casano, A., Verraes, A., Liem, R. K. H., Formstecher, E., Coppey-Moisan, M. and Galli, T. (2012). A Molecular Network for the Transport of the TI-VAMP/VAMP7 Vesicles from Cell Center to Periphery. Dev Cell 23, 166–180.

Burgo, A., Casano, A. M., Kuster, A., Arold, S. T., Wang, G., Nola, S., Verraes, A., Dingli, F., Loew, D. and Galli, T. (2013). Increased activity of the Vesicular Soluble N-Ethylmaleimide-sensitive Factor Attachment Protein Receptor TI-VAMP/VAMP7 by Tyrosine Phosphorylation in the Longin Domain. Journal of Biological Chemistry 288, 11960–11972.

Chaineau, M., Danglot, L., Proux-Gillardeaux, V. and Galli, T. (2008). Role of HRB in Clathrin-dependent Endocytosis. Journal of Biological Chemistry 283, 34365–34373.

Chaineau, M., Danglot, L. and Galli, T. (2009). Multiple roles of the vesicular-SNARE TI-VAMP in post-Golgi and endosomal trafficking. FEBS Lett 583, 3817–3826.

Charrin, S., le Naour, F., Silvie, O., Milhiet, P.-E., Boucheix, C. and Rubinstein, E. (2009). Lateral organization of membrane proteins: tetraspanins spin their web. Biochemical Journal 420, 133–154.

Chen, Y. A. and Scheller, R. H. (2001). SNARE-mediated membrane fusion. Nat Rev Mol Cell Biol 2, 98–106.

Chenouard, N., Smal, I., de Chaumont, F., Maška, M., Sbalzarini, I. F., Gong, Y., Cardinale, J., Carthel, C., Coraluppi, S., Winter, M., et al. (2014). Objective comparison of particle tracking methods. Nat Methods 11, 281–289.

Crescitelli, R., Lässer, C., Szabó, T. G., Kittel, A., Eldh, M., Dianzani, I., Buzás, E. I. and Lötvall, J. (2013). Distinct RNA profiles in subpopulations of extracellular vesicles: apoptotic bodies, microvesicles and exosomes. J Extracell Vesicles 2, 20677.

Crocker, J. C. and Grier, D. G. (1996). Methods of Digital Video Microscopy for Colloidal Studies. J Colloid Interface Sci 179, 298–310.

Danglot, L., Zylbersztejn, K., Petkovic, M., Gauberti, M., Meziane, H., Combe, R., Champy, M.-F., Birling, M.-C., Pavlovic, G., Bizot, J.-C., et al. (2012). Absence of TI-VAMP/Vamp7 Leads to Increased Anxiety in Mice. Journal of Neuroscience 32, 1962–1968.

Daste, F., Galli, T. and Tareste, D. (2015). Structure and function of longin SNAREs. J Cell Sci.

Diaz, E., Ayala, G., Diaz, M. E., Liang-Wei Gong and Toomre, D. (2010). Automatic Detection of Large Dense-Core Vesicles in Secretory Cells and Statistical Analysis of Their Intracellular Distribution. IEEE/ACM Trans Comput Biol Bioinform 7, 2–11.

Edgar, J. R., Manna, P. T., Nishimura, S., Banting, G. and Robinson, M. S. (2016). Tetherin is an exosomal tether. Elife 5,.

Escola, J.-M., Kleijmeer, M. J., Stoorvogel, W., Griffith, J. M., Yoshie, O. and Geuze, H. J. (1998). Selective Enrichment of Tetraspan Proteins on the Internal Vesicles of Multivesicular Endosomes and on Exosomes Secreted by Human B-lymphocytes. Journal of Biological Chemistry 273, 20121– 20127.

Fawcett, T. (2006). An introduction to ROC analysis. Pattern Recognit Lett 27, 861–874.

Galli, T., Zahraoui, A., Vaidyanathan, V. v., Raposo, G., Tian, J. M., Karin, M., Niemann, H. and Louvard, D. (1998). A Novel Tetanus Neurotoxin-insensitive Vesicle-associated Membrane Protein in SNARE Complexes of the Apical Plasma Membrane of Epithelial Cells. Mol Biol Cell 9, 1437– 1448.

Ge, S., Koseoglu, S. and Haynes, C. L. (2010). Bioanalytical tools for single-cell study of exocytosis. Anal Bioanal Chem 397, 3281–3304.

Guček, A., Gandasi, N. R., Omar-Hmeadi, M., Bakke, M., Døskeland, S. O., Tengholm, A. and Barg, S. (2019). Fusion pore regulation by cAMP/Epac2 controls cargo release during insulin exocytosis. Elife 8,.

Gundelfinger, E. D., Kessels, M. M. and Qualmann, B. (2003). Temporal and spatial coordination of exocytosis and endocytosis. Nat Rev Mol Cell Biol 4, 127–139.

Gupton, S. L. and Gertler, F. B. (2010). Integrin Signaling Switches the Cytoskeletal and Exocytic Machinery that Drives Neuritogenesis. Dev Cell 18, 725–736.

Han, J., Pluhackova, K. and Böckmann, R. A. (2017). The Multifaceted Role of SNARE Proteins in Membrane Fusion. Front Physiol 8,.

Hemler, M. E. (2005). Tetraspanin functions and associated microdomains. Nat Rev Mol Cell Biol 6, 801–811.

Hessvik, N. P., Sagini, K., Romero, S., Ramirez-Garrastacho, M., Rodriguez, M., Tutturen, A. E. V., Kvalvaag, A., Stang, E., Brech, A., Sandvig, K., et al. (2023). siRNA screening reveals that SNAP29 contributes to exosome release. Cellular and Molecular Life Sciences 80, 177.

Hu, Y., Osuna-Highley, E., Hua, J., Nowicki, T. S., Stolz, R., McKayle, C. and Murphy, R. F. (2010). Automated analysis of protein subcellular location in time series images. Bioinformatics 26, 1630– 1636.

Huang, S., Lifshitz, L. M., Jones, C., Bellve, K. D., Standley, C., Fonseca, S., Corvera, S., Fogarty, K. E. and Czech, M. P. (2007). Insulin Stimulates Membrane Fusion and GLUT4 Accumulation in Clathrin Coats on Adipocyte Plasma Membranes. Mol Cell Biol 27, 3456–3469.

Huang, X., Fan, J., Li, L., Liu, H., Wu, R., Wu, Y., Wei, L., Mao, H., Lal, A., Xi, P., et al. (2018). Fast, long-term, super-resolution imaging with Hessian structured illumination microscopy. Nat Biotechnol 36, 451–459.

Jahn, R. and Scheller, R. H. (2006). SNAREs — engines for membrane fusion. Nat Rev Mol Cell Biol 7, 631–643.

Jahn, R. and Südhof, T. C. (1999). Membrane Fusion and Exocytosis. Annu Rev Biochem 68, 863–911.

Jiang, S., Zhou, X., Kirchhausen, T. and Wong, S. T. C. (2007). Detection of molecular particles in live cells via machine learning. Cytometry Part A 71A, 563–575.

Johnson, G. R., Li, J., Shariff, A., Rohde, G. K. and Murphy, R. F. (2015). Automated Learning of Subcellular Variation among Punctate Protein Patterns and a Generative Model of Their Relation to Microtubules. PLoS Comput Biol 11, e1004614.

Jullie, D., Choquet, D. and Perrais, D. (2014). Recycling Endosomes Undergo Rapid Closure of a Fusion Pore on Exocytosis in Neuronal Dendrites. Journal of Neuroscience 34, 11106–11118.

Kowal, J., Arras, G., Colombo, M., Jouve, M., Morath, J. P., Primdal-Bengtson, B., Dingli, F., Loew, D., Tkach, M. and Théry, C. (2016). Proteomic comparison defines novel markers to characterize heterogeneous populations of extracellular vesicle subtypes. Proceedings of the National Academy of Sciences 113,.

Lagache, T., Grassart, A., Dallongeville, S., Faklaris, O., Sauvonnet, N., Dufour, A., Danglot, L. and Olivo-Marin, J.-C. (2018). Mapping molecular assemblies with fluorescence microscopy and object-based spatial statistics. Nat Commun 9, 698.

Lamichhane, T. N., Sokic, S., Schardt, J. S., Raiker, R. S., Lin, J. W. and Jay, S. M. (2015). Emerging Roles for Extracellular Vesicles in Tissue Engineering and Regenerative Medicine. Tissue Eng Part B Rev 21, 45–54.

Larios, J., Mercier, V., Roux, A. and Gruenberg, J. (2020). ALIX- and ESCRT-III–dependent sorting of tetraspanins to exosomes. Journal of Cell Biology 219,.

le Naour, F., André, M., Boucheix, C. and Rubinstein, E. (2006). Membrane microdomains and proteomics: Lessons from tetraspanin microdomains and comparison with lipid rafts. Proteomics 6, 6447–6454.

Levy, S. and Shoham, T. (2005). The tetraspanin web modulates immune-signalling complexes. Nat Rev Immunol 5, 136–148.

Li, D., Shao, L., Chen, B.-C., Zhang, X., Zhang, M., Moses, B., Milkie, D. E., Beach, J. R., Hammer, J. A., Pasham, M., et al. (2015). Extended-resolution structured illumination imaging of endocytic and cytoskeletal dynamics. Science (1979) 349,.

Lin, D., Lin, Z., Cao, J., Velmurugan, R., Ward, E. S. and Ober, R. J. (2019). A two-stage method for automated detection of ring-like endosomes in fluorescent microscopy images. PLoS One 14, e0218931.

Linkert, M., Rueden, C. T., Allan, C., Burel, J.-M., Moore, W., Patterson, A., Loranger, B., Moore, J., Neves, C., MacDonald, D., et al. (2010). Metadata matters: access to image data in the real world. Journal of Cell Biology 189, 777–782.

Liu, A., Huang, X., He, W., Xue, F., Yang, Y., Liu, J., Chen, L., Yuan, L. and Xu, P. (2021a). pHmScarlet is a pH-sensitive red fluorescent protein to monitor exocytosis docking and fusion steps. Nat Commun 12, 1413.

Liu, A., Huang, X., He, W., Xue, F., Yang, Y., Liu, J., Chen, L., Yuan, L. and Xu, P. (2021b). pHmScarlet is a pH-sensitive red fluorescent protein to monitor exocytosis docking and fusion steps. Nat Commun 12, 1413.

Mahmood, A., Otruba, Z., Weisgerber, A. W., Palay, M. D., Nguyen, M. T., Bills, B. L. and Knowles, M. K. (2023). Exosome secretion kinetics are controlled by temperature. Biophys J 122, 1301–1314.

Martineau, M., Somasundaram, A., Grimm, J. B., Gruber, T. D., Choquet, D., Taraska, J. W., Lavis, L. D. and Perrais, D. (2017). Semisynthetic fluorescent pH sensors for imaging exocytosis and endocytosis. Nat Commun 8, 1412.

Martinez-Arca, S., Alberts, P., Zahraoui, A., Louvard, D. and Galli, T. (2000). Role of Tetanus Neurotoxin Insensitive Vesicle-Associated Membrane Protein (Ti-Vamp) in Vesicular Transport Mediating Neurite Outgrowth. Journal of Cell Biology 149, 889–900.

Martinez-Arca, S., Coco, S., Mainguy, G., Schenk, U., Alberts, P., Bouillé, P., Mezzina, M., Prochiantz, A., Matteoli, M., Louvard, D., et al. (2001a). A Common Exocytotic Mechanism Mediates Axonal and Dendritic Outgrowth. The Journal of Neuroscience 21, 3830–3838.

Martinez-Arca, S., Coco, S., Mainguy, G., Schenk, U., Alberts, P., Bouillé, P., Mezzina, M., Prochiantz, A., Matteoli, M., Louvard, D., et al. (2001b). A Common Exocytotic Mechanism Mediates Axonal and Dendritic Outgrowth. The Journal of Neuroscience 21, 3830–3838.

Martinez-Arca, S., Rudge, R., Vacca, M., Raposo, G., Camonis, J., Proux-Gillardeaux, V., Daviet, L., Formstecher, E., Hamburger, A., Filippini, F., et al. (2003). A dual mechanism controlling the localization and function of exocytic v-SNAREs. Proceedings of the National Academy of Sciences 100, 9011–9016.

Mathieu, M., Névo, N., Jouve, M., Valenzuela, J. I., Maurin, M., Verweij, F. J., Palmulli, R., Lankar, D., Dingli, F., Loew, D., et al. (2021). Specificities of exosome versus small ectosome secretion revealed by live intracellular tracking of CD63 and CD9. Nat Commun 12, 4389.

Matsui, T., Osaki, F., Hiragi, S., Sakamaki, Y. and Fukuda, M. (2021). ALIX and ceramide differentially control polarized small extracellular vesicle release from epithelial cells. EMBO Rep 22,.

Mattheyses, A. L., Simon, S. M. and Rappoport, J. Z. (2010). Imaging with total internal reflection fluorescence microscopy for the cell biologist. J Cell Sci 123, 3621–3628.

Meldolesi, J. (2018). Exosomes and Ectosomes in Intercellular Communication. Current Biology 28, R435–R444.

Miesenböck, G., de Angelis, D. A. and Rothman, J. E. (1998). Visualizing secretion and synaptic transmission with pH-sensitive green fluorescent proteins. Nature 394, 192–195.

Miura, K. (2020). Bleach correction ImageJ plugin for compensating the photobleaching of time-lapse sequences. F1000Res 9, 1494.

Moro, A., Hoogstraaten, R. I., Persoon, C. M., Verhage, M. and Toonen, R. F. (2021a). Quantitative analysis of dense-core vesicle fusion in rodent CNS neurons. STAR Protoc 2, 100325.

Moro, A., van Nifterick, A., Toonen, R. F. and Verhage, M. (2021b). Dynamin controls neuropeptide secretion by organizing dense-core vesicle fusion sites. Sci Adv 7,.

Olivo-Marin, J.-C. (2002). Extraction of spots in biological images using multiscale products. Pattern Recognit 35, 1989–1996.

Persoon, C. M., Hoogstraaten, R. I., Nassal, J. P., van Weering, J. R. T., Kaeser, P. S., Toonen, R. F. and Verhage, M. (2019). The RAB3-RIM Pathway Is Essential for the Release of Neuromodulators. Neuron 104, 1065–1080.e12.

Proux-Gillardeaux, V., Rudge, R. and Galli, T. (2005). The Tetanus Neurotoxin-Sensitive and Insensitive Routes to and from the Plasma Membrane: Fast and Slow Pathways? Traffic 6, 366–373.

Proux-Gillardeaux, V., Raposo, G., Irinopoulou, T. and Galli, T. (2007). Expression of the Longin domain of TI-VAMP impairs lysosomal secretion and epithelial cell migration. Biol Cell 99, 261– 271.

Püspöki, Z., Sage, D., Ward, J. P. and Unser, M. (2016). SpotCaliper: fast wavelet-based spot detection with accurate size estimation. Bioinformatics 32, 1278–1280.

Roizin, L., Nishikawa, K., Koizumi, J. and Keoseian, S. (1967). The Fine Structure of the Multivesicular Body and Their Relationship to the Ultracellular Constituents of the Central Nervous System * ^†^. J Neuropathol Exp Neurol 26, 223–249.

Roth, J., Mehl, J. and Rohrbach, A. (2020). Fast TIRF-SIM imaging of dynamic, low-fluorescent biological samples. Biomed Opt Express 11, 4008.

Rubinstein, E., Le Naour, F., Lagaudrière-Gesbert, C., Billard, M., Conjeaud, H. and Boucheix, C. (1996). CD9, CD63, CD81, and CD82 are components of a surface tetraspan network connected to HLA-DR and VLA integrins. Eur J Immunol 26, 2657–2665.

Ruusuvuori, P., Äijö, T., Chowdhury, S., Garmendia-Torres, C., Selinummi, J., Birbaumer, M., Dudley, A. M., Pelkmans, L. and Yli-Harja, O. (2010). Evaluation of methods for detection of fluorescence labeled subcellular objects in microscope images. BMC Bioinformatics 11, 248.

Sankaranarayanan, S., de Angelis, D., Rothman, J. E. and Ryan, T. A. (2000a). The Use of pHluorins for Optical Measurements of Presynaptic Activity. Biophys J 79, 2199–2208.

Sankaranarayanan, S., De Angelis, D., Rothman, J. E. and Ryan, T. A. (2000b). The Use of pHluorins for Optical Measurements of Presynaptic Activity. Biophys J 79, 2199–2208.

Schindelin, J., Arganda-Carreras, I., Frise, E., Kaynig, V., Longair, M., Pietzsch, T., Preibisch, S., Rueden, C., Saalfeld, S., Schmid, B., et al. (2012). Fiji: an open-source platform for biological-image analysis. Nat Methods 9, 676–682.

Schmoranzer, J., Goulian, M., Axelrod, D. and Simon, S. M. (2000). Imaging Constitutive Exocytosis with Total Internal Reflection Fluorescence Microscopy. Journal of Cell Biology 149, 23–32.

Sebastian, R., Diaz, M.-E., Ayala, G., Letinic, K., Moncho-Bogani, J. and Toomre, D. (2006). Spatio-Temporal Analysis of Constitutive Exocytosis in Epithelial Cells. IEEE/ACM Trans Comput Biol Bioinform 3, 17–32.

Shen, Y., Rosendale, M., Campbell, R. E. and Perrais, D. (2014). pHuji, a pH-sensitive red fluorescent protein for imaging of exo- and endocytosis. Journal of Cell Biology 207, 419–432.

Smal, I., Loog, M., Niessen, W. and Meijering, E. (2010). Quantitative Comparison of Spot Detection Methods in Fluorescence Microscopy. IEEE Trans Med Imaging 29, 282–301.

Théry, C., Witwer, K. W., Aikawa, E., Alcaraz, M. J., Anderson, J. D., Andriantsitohaina, R., Antoniou, A., Arab, T., Archer, F., Atkin-Smith, G. K., et al. (2018). Minimal information for studies of extracellular vesicles 2018 (MISEV2018): a position statement of the International Society for Extracellular Vesicles and update of the MISEV2014 guidelines. J Extracell Vesicles 7, 1535750.

Toonen, R. F., Kochubey, O., de Wit, H., Gulyas-Kovacs, A., Konijnenburg, B., Sørensen, J. B., Klingauf, J. and Verhage, M. (2006). Dissecting docking and tethering of secretory vesicles at the target membrane. EMBO J 25, 3725–3737.

Urbina, F. and Gupton, S. L. (2021). Automated Detection and Analysis of Exocytosis. Journal of Visualized Experiments.

Urbina, F. L., Gomez, S. M. and Gupton, S. L. (2018). Spatiotemporal organization of exocytosis emerges during neuronal shape change. Journal of Cell Biology 217, 1113–1128.

van Deventer, S., Arp, A. B. and van Spriel, A. B. (2021). Dynamic Plasma Membrane Organization: A Complex Symphony. Trends Cell Biol 31, 119–129.

van Niel, G., D’Angelo, G. and Raposo, G. (2018). Shedding light on the cell biology of extracellular vesicles. Nat Rev Mol Cell Biol 19, 213–228.

Vats, S. and Galli, T. (2022). Role of SNAREs in Unconventional Secretion—Focus on the VAMP7-Dependent Secretion. Front Cell Dev Biol 10,.

Verderio, C., Cagnoli, C., Bergami, M., Francolini, M., Schenk, U., Colombo, A., Riganti, L., Frassoni, C., Zuccaro, E., Danglot, L., et al. (2012). TI-VAMP/VAMP7 is the SNARE of secretory lysosomes contributing to ATP secretion from astrocytes. Biol Cell 104, 213–228.

Verhage, M. and Sørensen, J. B. (2008). Vesicle Docking in Regulated Exocytosis. Traffic 9, 1414– 1424.

Verweij, F. J., Bebelman, M. P., Jimenez, C. R., Garcia-Vallejo, J. J., Janssen, H., Neefjes, J., Knol, J. C., de Goeij-de Haas, R., Piersma, S. R., Baglio, S. R., et al. (2018). Quantifying exosome secretion from single cells reveals a modulatory role for GPCR signaling. Journal of Cell Biology 217, 1129–1142.

Verweij, F. J., Bebelman, M. P., George, A. E., Couty, M., Bécot, A., Palmulli, R., Heiligenstein, X., Sirés-Campos, J., Raposo, G., Pegtel, D. M., et al. (2022). ER membrane contact sites support endosomal small GTPase conversion for exosome secretion. Journal of Cell Biology 221,.

Wang, G., Nola, S., Bovio, S., Bun, P., Coppey-Moisan, M., Lafont, F. and Galli, T. (2018). Biomechanical Control of Lysosomal Secretion Via the VAMP7 Hub: A Tug-of-War between VARP and LRRK1. iScience 4, 127–143.

Witwer, K. W. and Théry, C. (2019). Extracellular vesicles or exosomes? On primacy, precision, and popularity influencing a choice of nomenclature. J Extracell Vesicles 8, 1648167.

Wu, L.-G., Hamid, E., Shin, W. and Chiang, H.-C. (2014). Exocytosis and Endocytosis: Modes, Functions, and Coupling Mechanisms. Annu Rev Physiol 76, 301–331.

Yáñez-Mó, M., Siljander, P. R.-M., Andreu, Z., Bedina Zavec, A., Borràs, F. E., Buzas, E. I., Buzas, K., Casal, E., Cappello, F., Carvalho, J., et al. (2015). Biological properties of extracellular vesicles and their physiological functions. J Extracell Vesicles 4, 27066.

Yuan, T., Lu, J., Zhang, J., Zhang, Y. and Chen, L. (2015). Spatiotemporal Detection and Analysis of Exocytosis Reveal Fusion “Hotspots” Organized by the Cytoskeleton in Endocrine Cells. Biophys J 108, 251–260.

